# Development of a fixed list of descriptors for the qualitative behavioural assessment of shelter dogs

**DOI:** 10.1101/545020

**Authors:** Laura Arena, Françoise Wemelsfelder, Stefano Messori, Nicola Ferri, Shanis Barnard

## Abstract

The shelter environment may have a severe impact on the quality of life of dogs, and there is thus a need to develop valid tools to assess their welfare. These tools should be sensitive not only to the animals’ physical health but also to their mental health, including the assessment of positive and negative emotions. Qualitative Behaviour Assessment (QBA) is an integrative ‘whole animal’ measure that captures the expressive quality of an animal’s demeanour, using descriptors such as ‘relaxed’, ‘anxious’, and ‘playful’. In this study, for the first time, we developed and tested a fixed-list of qualitative QBA descriptors for application to dogs living in kennels. A list of 20 QBA descriptors was developed based on literature search and an expert opinion survey. Inter-observer reliability was investigated by asking 11 observers to use these descriptors to score 13 video clips of kennelled dogs. Principal Component Analysis (PCA) was used to extract four main dimensions together explaining 70.9% of the total variation between clips. PC1 characterised curious/playful/excitable, sociable demeanour, PC2 ranged from comfortable/relaxed to anxious/nervous/stressed expression, PC3 described fearful demeanour, and PC4 characterized bored/depressed demeanour. Observers’ agreement on the ranking of video clips on these four expressive dimensions was good (Kendall’s W: 0.60-0.80). ANOVA showed a significant effect of observer on mean clip score on all PCs (p<0.05) due to a few observers scoring differently from the rest of the group. These results indicate the potential of the proposed list of QBA terms for sheltered dogs to serve as a non-invasive, easy-to-use assessment tool. However, the observers’ effect on mean scores points towards the need for adequate observer training. The QBA scoring tool can be integrated with existing welfare assessment protocols for shelter dogs and strengthen the power of those protocols to assess and evaluate the animals’ experience in shelters.

## 1. INTRODUCTION

Kennel dog welfare is a concern affecting thousands of animals all around the world held in temporary or permanent confinement for a variety of reasons [1,2]. There is evidence that shelter environments may have a severe impact on the quality of life of dogs [3,4]. This is likely due to factors such as social isolation and novel surroundings [5], especially if these are protracted over long periods of time [6]. For this reason, there has been an increasing interest by the scientific community to develop validated tools to assess the welfare of sheltered dogs [3,7].

Over the last decades, the concept of animal welfare has evolved from focusing primarily on the animal’s physical health and ability to cope with its environment [8], to recognising that animals are sentient beings capable of experiencing positive and negative emotions [9]. It is now accepted that animals, though healthy, can nevertheless experience negative emotions when housed in unsuitable environments [10]. In addition the concept of animal welfare has shifted in recent times from focusing on the reduction of negative emotions (e.g. fear, pain) to ensuring that captive/domestic animals are also experiencing positive emotions (e.g. pleasure, happiness) [11,12]. It is therefore essential that animal welfare assessment tools include measures of positive welfare.

The potential of qualitative methods for assessing behaviour and welfare to play a role in such developments, and, alongside quantitative measure, to contribute to useful measures of positive animal welfare, has been a subject of review and discussion [13,14]. Qualitative Behavioural Assessment (QBA) is one such method which to date has generated a substantial body of research assessing emotional expressivity in a range of animal species [15–18]. QBA is a ‘whole-animal’ approach measuring not so much *what* the animal is doing physically, as *how* it is performing these behaviours, i.e. the expressive style or demeanour with which the animal is moving [19]. QBA uses a range of positive and negative qualitative descriptors such as ‘relaxed’, ‘sociable’, ‘anxious’ or ‘fearful’ in order to quantify an animal’s experience in different situations and conditions [19].

The descriptive terms used in QBA can either be developed through an experimental procedure known as Free-Choice Profiling (FCP), in which each assessor generates his/her own descriptors based on the observation of animals in different situations [20,21], or are provided in the form of a pre-determined list of QBA descriptors given to observers to assess animals. When standardisation of measurement tools is required, for example for the purpose of on-farm welfare inspections, the use of a fixed-list designed for the purpose of welfare assessment in a particular species is more feasible than FCP. Pre-fixed QBA protocols are for example included in Welfare Quality^®^ protocols for cattle, poultry and pigs [22] and in AWIN protocols for sheep, goats, horses, and donkeys [23–25], where they serve as a measure for positive emotional state.

Initial validation of the application of QBA to dogs was done by Walker et al. [26,27]. In these two studies Walker and colleagues applied FCP for the first time to assessing emotional expressivity in working dogs (all Beagles), and in shelter dogs in home pen and novel test pen environments. Outcomes showed, in the different dog groups, overlapping dimensions of emotional expressivity such as confidence/contentment, anxiousness/unsureness, alertness/attentiveness or curiousness/inquisitiveness. Walker et al. [27] also showed significant and meaningful correlations between QBA dimensions and quantitative ethogram-based behavioural measures, suggesting the potential value of QBA as a welfare assessment tool for dogs. Arena and colleagues [28] further investigated this potential by applying FCP to shelter dogs in a wide variety of shelter environments and social contexts including outdoor/indoor pens, single/pair/group housing, and the presence/absence of human activity. Results showed that by presenting more complex contexts and giving the animals more opportunities to express a wider repertoire of emotions, the observers generated ‘richer’ expressive dimensions than was the case in Walker et al.’s [26,27] studies which used relatively standardised experimental settings.

The present study aimed to build on these outcomes by developing a fixed-term QBA descriptor list that might be integrated with existing on-shelter welfare assessment protocols. Adding QBA could provide information complementary to that provided by quantitative measures, extending a protocol’s power to identify and detect emotional shifts in dogs across the positive and negative emotional spectrum [7,27,28]. Recognising that QBA relies on context-specific qualitative judgment of behavioural expression [19], descriptors included in a pre-fixed list should be representative of the large range of behavioural expressions that dogs could potentially show in variable kennel conditions. To achieve this goal, we generated a list of suitable terms based on the available dog behaviour and welfare literature, and then used an expert opinion survey to refine this list into a final set of 20 QBA terms. The inter-observer reliability of these terms was subsequently investigated by instructing eleven observers to score the emotional expressivity of dogs viewed in a sample of videos reflecting a wide range of shelter conditions.

## 2. MATERIAL AND METHODS

### 2.1 Ethics statement

No special permission for use of dogs in such behavioural studies is required in Italy, since dogs were observed during their daily life and within their familiar shelter pens. When dogs were exposed to people, no physical interaction was required.

All procedures were performed in full accordance with Italian legal regulations and the guidelines for the treatments of animals in behavioural research and teaching of the Association for the Study of Animal Behavior (ASAB).

No IRB approval was sought for the use of students as observers, but they provided informed signed consent to participate in the study and they were fully informed about the purpose and background of the study. Students ranged in age from 25 and 34 years.

### 2.2 Selection of terms

#### 2.2.1 Literature review

In a previous QBA study by this research group [28], 13 observers generated a number of terms to describe the emotional expression of shelter dogs from videos, using a Free Choice Profiling (FCP) methodology [15]. The FCP in Arena et al. [28] generated over 50 terms and the analysis extracted three main consensus dimensions. From that work, we selected 16 high-loading terms characterising each of the three main dimensions (Table 1). These 16 terms were used as a starting-point for the current study. Additional qualitative descriptors were selected from the relevant scientific literature on dogs’ emotions [26,27,29], behaviour [30], personality and temperament [31–34], and from other QBA research [13,35,36]. Thus, nine new terms were added (affectionate, aloof, attention-seeking, boisterous, excited, explorative, happy, self-confident and serene), creating a preliminary list of 25 terms.

**Table 1:** List of terms summarising the highest positive and negative loadings on each consensus dimension generated through Free Choice Profile methodology in Arena et al. [28].

|  | Positive | Negative |
| --- | --- | --- |
| Dimension 1 | playful/sociable/curious | bored/uncomfortable/apathetic |
| Dimension 2 | relaxed/tranquil | nervous/alert/fearful |
| Dimension 3 | stressed/bored/anxious | wary/timorous/hesitant |

*2.2.2 Expert opinion*

#### 2.2.2 Expert opinion

To check for the appropriateness, relevance and ease of understanding of the 25 terms on the preliminary list, we performed an expert opinion survey. For this purpose, we selected a panel of 15 international experts in the field of dog personality and behaviour, shelter dog welfare and QBA methodology. The panel was composed of 12 females and three males. A one-round on-line survey was arranged using the SurveyMonkey^®^ platform. In an introductory letter, experts were told that the goal of the project was to develop a comprehensive list of terms for qualitative assessment of the emotional state and welfare of dogs housed in different shelter environments. Eight out of 15 experts answered the survey anonymously.

For each descriptor, we provided a brief semantic characterisation and asked the experts to score that term on the basis of four brief statements using a likert-scale from 1 to 5, where 1 corresponded to ‘not at all’, and 5 to ‘completely’ (Table 2). Experts also had the possibility to add comments in a free space if wanted. Furthermore, at the end of the survey the experts could suggest a maximum of three terms that, in their opinion, were missing but that they considered important to describe the emotional state of sheltered dogs. This expert opinion was used to refine our original list by deleting and/or adding terms, in order to eliminate synonyms, select the terms most relevant to describing shelter dog emotional state, and ensure that terms were easy to understand.

**Table 2.** Mean scores for the four statements presented in the survey for each term.

| In your opinion, this descriptor is: | Survey mean scores* |
| --- | --- |
| 1. easily comprehensible and intuitive | 3.60 |
| 2. ambiguous, might be interpreted with different meanings | 2.75 |
| 3. useful to describe the emotional state of sheltered dogs | 3.49 |
| 4. relevant to assess the welfare of sheltered dogs | 3.25 |

\*calculated by averaging for each statement the experts’ scores across all terms.
*2.2.3 Generating the final list*

#### 2.2.3 Generating the final list

The SurveyMonkey^®^ software automatically generated a matrix with the average scores assigned by the experts to each term. We then calculated the mean scores of each of the four statements for all terms (Table 2). To decide whether to keep or delete a term, we used those total mean scores to establish thresholds for each statement: to be considered for inclusion, a term had to receive a mean score of 3 or more in statements #1, #3 and #4, and less than 3 in statement #2. Furthermore, we summarised and classified the negative comments provided by the experts as ‘prone to anthropomorphism’; ‘too generic’; ‘too similar to other terms’; ‘is not an emotional state’. We put aside inapplicable comments such as those repeating the same concept expressed in the statement (e.g. “not useful to assess animal welfare”), those referring to the definition given by the authors (e.g. “I would include ‘interest’ in the definition”) and comments without justification (e.g. “I suggest to delete this term from the list”).

Finally, we defined inclusion/exclusion criteria in order to refine our list.

As exclusion criteria, we established that a descriptor would be excluded from the list if it had at least:

– two insufficient scores
– one insufficient score and one negative comment
– no insufficient scores, but three or more negative comments.

### 2.3 Inter-observer reliability

The final list of terms resulting from the previous steps was subsequently used to test for inter-observer reliability.

#### 2.3.1 Animals and video-clips

A convenience sample of dogs in thirteen kennels were video-recorded in seven different Italian shelters: three in Northern Italy (Emilia-Romagna and Piemonte Regions), two in the Centre (Abruzzo Region) and two in the South (Puglia Region). The aim was to record dogs in different environments and scenarios to capture a variety of dog behavioural expressions. All dogs were in the kennels on a long-term basis (> 6 months since admittance). In each kennel, all resident dogs were video-recorded using a smartphone (Samsung GT-I9100P) that had been mounted on a tripod positioned a few meters from the fence. Dogs were housed singly in 4 videos, in pairs in another 4 videos and in groups of 3 or more dogs in 5 videos. Nine kennels were located outdoors and four indoors.

To trigger different responses, dogs were recorded during three different scenarios common in a shelter environment: under normal conditions with no external intervention, in the presence of an unknown person, and in presence of a familiar person. The unfamiliar person was one of two researchers (one female and one male), while the familiar person was a shelter operator available at the time of recording. The unfamiliar person followed a simple protocol, approaching the fence of the kennel and standing one metre from the fence ignoring the dogs (1 min) and subsequently talking gently to the dog moving a hand slowly along the fence (1 min). Shelter operators were asked to enter the kennel and interact with the dog/s for 2 minutes.

The 13 recorded clips were cut (using the Avidemux 2.6.8 programme) to obtain clips of 1.5 minutes average length during which all dog/s in the kennel were visible at all time.

#### 2.3.2 Observers and observation session

Eleven students enrolled in a course for dog trainers were recruited as observers (8 females and 3 males). All participants had previous experience with dogs and owned at least one dog. Five of them were dog trainers, four were volunteers at dog shelters, two had graduated at the Faculty of Animal Welfare and Protection, and one was a veterinarian. None of the observers had previous experience with QBA.

Before starting the assessment of the video clips, approximately 1 hour was dedicated to an introduction with the goal of explaining the aim of the study and the operative procedures. A brief characterisation of each term was provided and, where necessary, terms were discussed further among the observers.

Observers were told that the study had the aim of investigating whether observers can agree in using a fixed list of QBA terms to assess emotional expressivity in shelter dogs. Emotional expressivity was defined as an animal’s style of interaction with its environment, conspecifics and humans (how the animal behaves as opposed to what it does). Each observer was provided with scoring sheets (one for each video clip) on which Visual Analogue Scales (VAS) of 125 mm of length were placed next to each term. Participants were told to use every qualitative descriptor in the list to score the animals visible in the clips. They were instructed on how to use the VAS for scoring: the left end of the scale corresponded to the minimum score (0 mm), meaning the expressive quality indicated by the term was entirely absent in that dog or group of dogs, whereas the right end represented the maximum score (125 mm), meaning that the expressive quality indicated by the term was strongly dominant in that dog or group of dogs. Observers were asked to avoid talking during the session.

Once instructions had ended, observers watched the 13 clips projected onto a lecture hall screen and, after each clip, they had approximately 2 mins to score the animals’ expressions on the rating scales, by drawing a vertical line across the VAS at the point they felt was appropriate. A score was assigned to each term for each clip by measuring with a ruler the distance in millimetres between the minimum point of the VAS and the point where the observer marked the line.

#### 2.3.3 Statistical analysis

IBM SPSS Statistics 21 software (IBM Corp., 2012) was used for statistical analysis. QBA data were analysed using a Principal Component Analysis (PCA, correlation matrix, varimax rotation). Before running the PCA, we explored the sampling adequacy by checking Kaiser-Meyer-Olkin (KMO) and anti-correlation image matrix values. The output scores of the main dimensions extracted by the PCA were then used to test the inter-observer reliability, using Kendall’s W coefficient of concordance [37]. Kendall’s W values can vary from 0 (no agreement at all) to 1 (complete agreement), with values higher than 0.6 showing substantial agreement. To check that mean scores did not differ between observers, a one-way ANOVA was used on each of the four PCA dimensions with observer as fixed factor and video clip as random factor. Post-hoc multiple comparisons (using Tukey HSD test) and homogeneous subsets were inspected to check which and how many observers were in disagreement. Finally, we assessed the inter-observer agreement for each descriptor separately, again using the Kendall’s W test.

## 3. RESULTS

### 3.1 Generation of the list of terms

Mean scores attributed to each statement and for each term by the panel of responding experts are summarised in Table 3.

**Table 3.** Mean scores attributed by the experts to each term for each statement. <u>Underlined</u> are the ‘insufficient’ scores and in **bold** the deleted terms. #Q: numbers 1 to 4 correspond to the statements as listed in Table 2.

| Terms | #Q | Score | Terms | #Q | Score | Terms | #Q | Score |
| --- | --- | --- | --- | --- | --- | --- | --- | --- |
| Playful | 1 | 4.38 | Nervous | 1 | 3.63 | Wary | 1 | 3.75 |
|  | 2 | 1.75 |  | 2 | 2.75 |  | 2 | 3.00 |
|  | 3 | 4.13 |  | 3 | 3.75 |  | 3 | 4.13 |
|  | 4 | 4.13 |  | 4 | 3.13 |  | 4 | 3.88 |
| Sociable | 1 | 4.25 | Alert | 1 | 3.75 | Hesitant | 1 | 3.88 |
|  | 2 | 1.88 |  | 2 | 2.75 |  | 2 | 2.75 |
|  | 3 | 4.00 |  | 3 | 4.00 |  | 3 | 3.00 |
|  | 4 | 4.13 |  | 4 | 3.00 |  | 4 | <u>2.75</u> |
| Curious | 1 | 3.75 | **Boisterous** | 1 | <u>2.38</u> | **Aloof** | 1 | 3.00 |
|  | 2 | 2.00 |  | 2 | <u>3.75</u> |  | 2 | <u>3.13</u> |
|  | 3 | 3.50 |  | 3 | <u>2.88</u> |  | 3 | <u>2.63</u> |
|  | 4 | 3.25 |  | 4 | <u>2.50</u> |  | 4 | <u>2.75</u> |
| **Happy** | 1 | 3.88 | Excited | 1 | 3.25 | Fearful | 1 | 4.75 |
|  | 2 | 2.75 |  | 2 | 2.63 |  | 2 | 1.63 |
|  | 3 | 3.50 |  | 3 | 3.75 |  | 3 | 4.50 |
|  | 4 | 3.25 |  | 4 | 3.00 |  | 4 | 4.25 |
| **Affectionate** | 1 | 3.25 | **Apathetic** | 1 | 3.13 | **Timorous** | 1 | 3.25 |
|  | 2 | 2.13 |  | 2 | <u>3.13</u> |  | 2 | <u>3.25</u> |
|  | 3 | 3.38 |  | 3 | <u>2.88</u> |  | 3 | 3.38 |
|  | 4 | <u>2.88</u> |  | 4 | <u>2.63</u> |  | 4 | <u>2.88</u> |
| Attention-seeking | 1 | 3.88 | Uncomfortable | 1 | 3.25 | Stressed | 1 | 3.88 |
|  | 2 | 2.38 |  | 2 | 2.75 |  | 2 | 2.38 |
|  | 3 | 3.38 |  | 3 | 3.25 |  | 3 | 4.38 |
|  | 4 | 3.25 |  | 4 | 3.50 |  | 4 | 4.63 |
| Relaxed | 1 | 3.50 | Bored | 1 | 3.25 | Anxious | 1 | 4.25 |
|  | 2 | 2.88 |  | 2 | 2.75 |  | 2 | 1.88 |
|  | 3 | 3.50 |  | 3 | 3.50 |  | 3 | 4.50 |
|  | 4 | 3.25 |  | 4 | 3.50 |  | 4 | 4.50 |
| Tranquil | 1 | <u>2.50</u> | Self-confident | 1 | 3.50 | Explorative | 1 | 3.88 |
|  | 2 | <u>3.25</u> |  | 2 | 2.88 |  | 2 | 2.00 |
|  | 3 | <u>2.88</u> |  | 3 | 3.00 |  | 3 | 3.25 |
|  | 4 | <u>2.63</u> |  | 4 | <u>2.88</u> |  | 4 | 3.38 |
| Serene | 1 | <u>2.88</u> |  |  |  |  |  |  |
|  | 2 | <u>3.50</u> |  |  |  |  |  |  |
|  | 3 | <u>2.00</u> |  |  |  |  |  |  |
|  | 4 | <u>2.00</u> |  |  |  |  |  |  |

On the basis of the inclusion/exclusion criteria, nine terms were deleted from the list: boisterous, aloof, timorous, tranquil, serene, apathetic (all had at least two insufficient scores, and some also had negative comments), affectionate, self-confident (one insufficient score and at least one negative comment), happy (at least three negative comments). We evaluated the additional terms suggested by the panel of experts, and we added interested, depressed and aggressive (each one, suggested three times) and reactive (suggested twice). Finally, the term uncomfortable was replaced by the positive form comfortable.

As a result, the final list was composed of 20 terms. (Table 4). Overall this list was balanced in term of their semantic meaning (*positive* versus *negative* emotional state) and level of arousal they expressed (*high* versus *low* arousal). A brief characterisation of each term was produced on the basis of previous QBA studies [25–27]. These characterisations were then translated onto Italian to facilitate scoring by the observers.

**Table 4.**
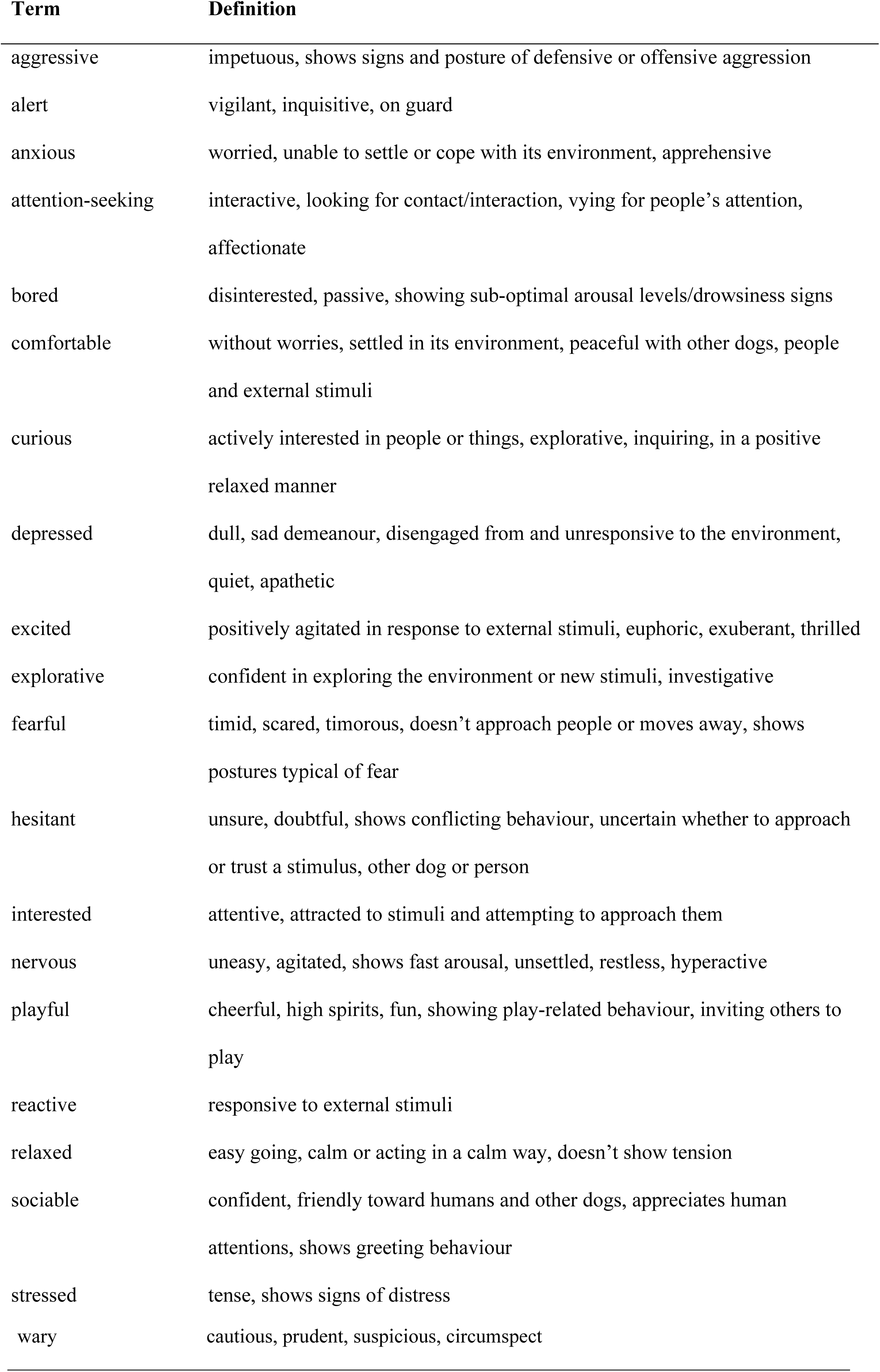
Final list of terms and their characterisations

### Inter-observer reliability of terms

KMO was well above the threshold of 0.50 (0.83) and the anti-image correlations were also high for all individual variables (> 0.55). After visual inspection of the scree plot, four components were extracted which together explained 70.9% of the total variance and all with Eigenvalues >1 (Table 5). The observers’ agreement on these four dimensions varied between 0.60 and 0.80, all significant at *p* < 0.001 (Table 5).

**Table 5.** Principal component (PC) analysis outcomes and inter-observer agreement (using Kendall’s W) for QBA rating scales.

|  | PC1 | PC2 | PC3 | PC4 |
| --- | --- | --- | --- | --- |
| **Eigenvalue** | 5.66 | 5.18 | 1.78 | 1.56 |
| **% explained variance** | 28.3 | 25.9 | 8.9 | 7.8 |
| **% cumulative variance** | 28.3 | 54.2 | 63.1 | 70.9 |
| **Kendall W** | 0.61 | 0.80 | 0.71 | 0.60 |
| **p-value** | $p < 0.001$ | $p < 0.001$ | $p < 0.001$ | $p < 0.001$ |

Table 6 shows the loadings of each descriptor on the four principal components (PC). PC1 was characterized by positive terms curious/attention-seeking/playful/excited/sociable/interested and explorative; PC2 characterized dogs as ranging from comfortable/relaxed to anxious/nervous/stressed. Figure 1 shows the 20 QBA descriptors plotted along these first two PCs. PC3 was characterized by the terms fearful/hesitant/wary, and finally, PC4 by the terms depressed/bored. Figure 2 shows the 20 QBA descriptors plotted along PC3 and PC4.

**Table 6.**
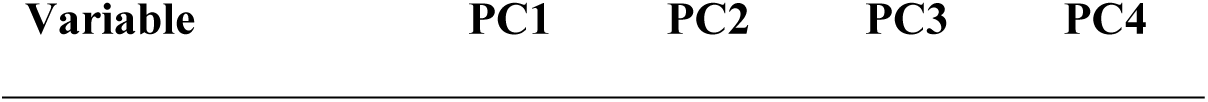
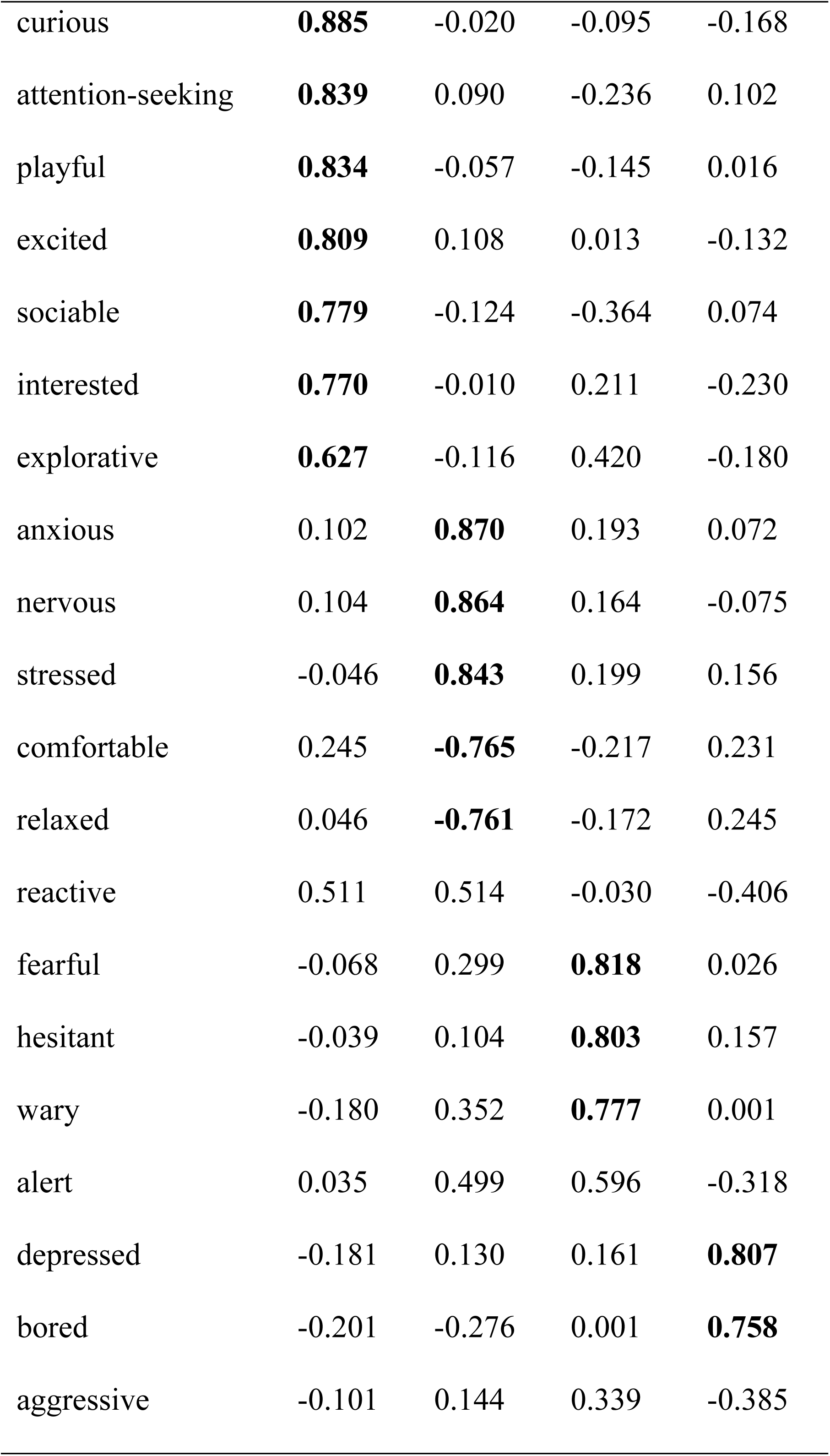
Loadings for each QBA descriptor on the four principal components (PC1, PC2, PC3 and PC4) extracted by the PCA analysis. Loadings higher than 0.60 for each term are bold typed.

**Figure 1.**
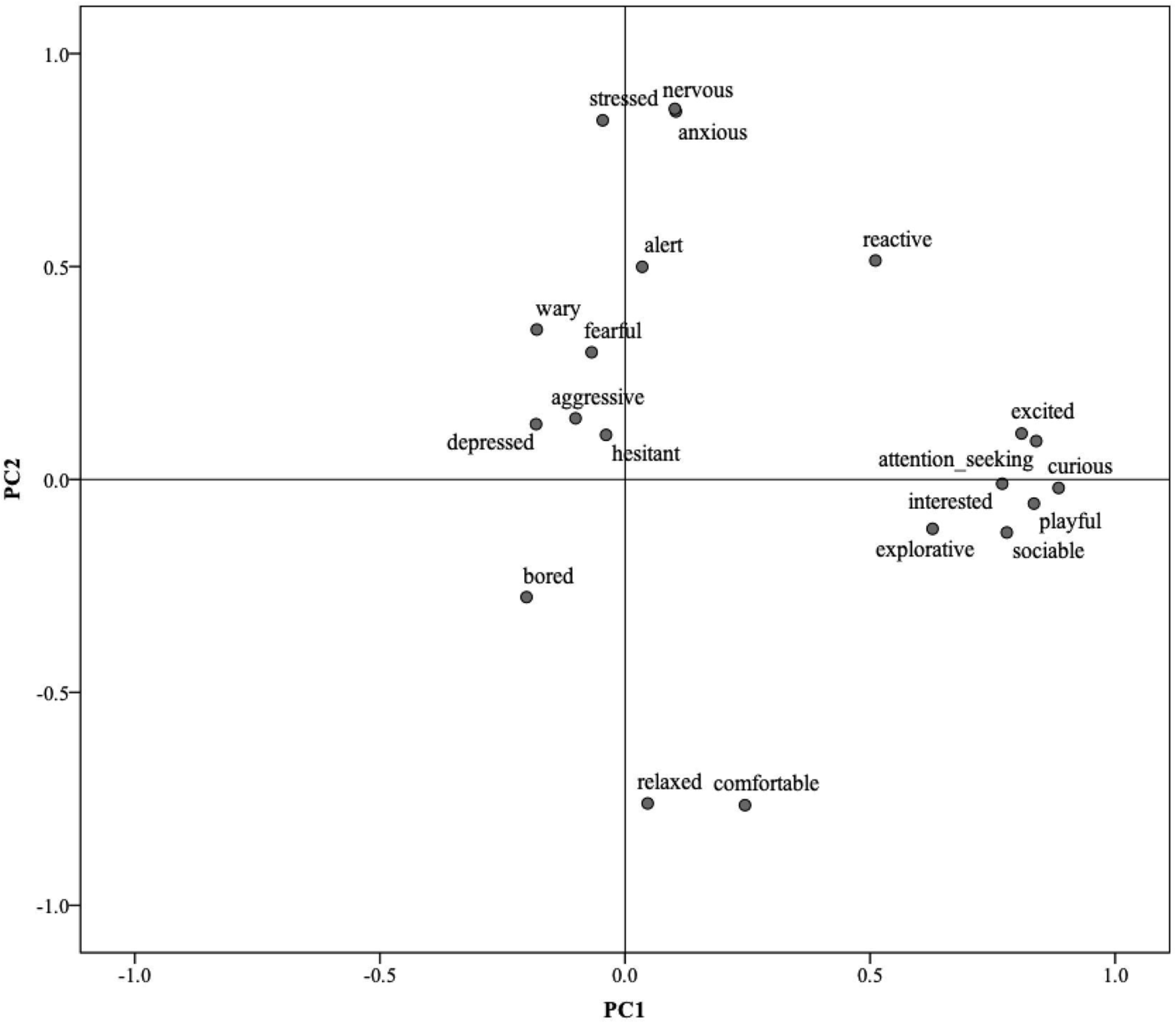
Distribution of the 20 QBA descriptors on Component 1 (PC1) and Component 2 (PC2) plot rotated in space.

**Figure 2.**
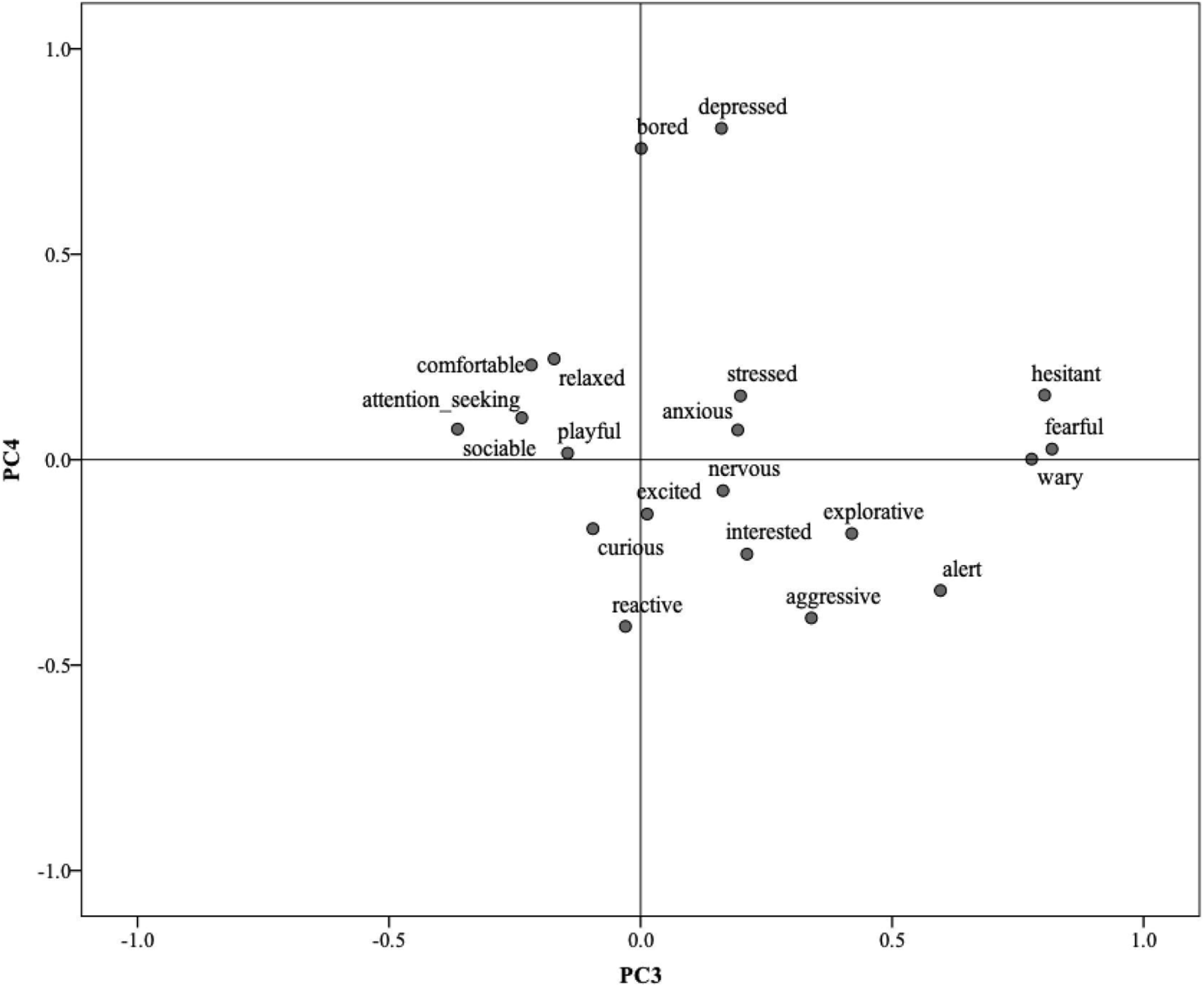
Distribution of the 20 QBA descriptors on Component 3 (PC3) and Component 4 (PC4) plot rotated in space.

ANOVA showed that there was a significant effect of observer on mean QBA scores on all four PCs (PC1: F = 13.7, p < 0.001; PC2: F = 3.5, p < 0.001; PC3: F=15.6, p<0.001; PC4: F=6.4, p<0.001). This means that although observers were in agreement on the ranking of the dogs in the videos, their average clip scores for the different PCs differed. Inspection of post-hoc multiple comparisons indicated that for PCs 1, 3 and 4 most observers (>7) presented homogeneous means, with the remaining 3-4 observers scoring differently from the rest of the group. The observers scoring differently were often the same across all PCs. Post-hoc analysis for PC2 highlighted one clear outlier (observer 10), with the remaining observers not differing from each other in mean clip score. When running the ANOVA again, without observer 10, observer effect was not significant (F=1.62, p=0.12).

The Kendall W values for individual descriptors were all significantly different from chance (p<0.001), but only 8 terms reached values higher than 0.60, while the term ‘depressed’ showed a particularly low value (Table 7).

**Table 7.** Kendall’s W coefficients of concordance (n=11) for the 20 used QBA descriptors. Values larger than 0.60 are bold typed.

| Variable | Kendall W | Variable | Kendall W |
| --- | --- | --- | --- |
| Aggressive | 0.489 | Fearful | 0.561 |
| Alert | 0.579 | Hesitant | 0.524 |
| Anxious | **0.686** | Interested | 0.503 |
| Attention-seeking | **0.679** | Nervous | **0.683** |
| Bored | 0.540 | Playful | 0.563 |
| Comfortable | **0.700** | Reactive | **0.611** |
| Curious | 0.576 | Relaxed | **0.689** |
| Depressed | 0.320 | Sociable | **0.793** |
| Excited | 0.497 | Stressed | 0.579 |
| Explorative | 0.466 | Wary | **0.714** |

## 4. DISCUSSION

The aim of this research was to develop and test the inter-observer reliability of a fixed Qualitative Behaviour Assessment (QBA) rating scale for the purpose of field assessment of emotional expressivity in dogs in a shelter environment. Attempts had previously been made to develop and validate welfare assessment protocols for kennelled dogs [4,7], however, these protocols did not include measures addressing animals’ emotions. To our knowledge, our study is the first to generate a fixed QBA rating scale for the assessment of shelter dogs’ emotion. A previous Free Choice Profiling study served as a basis for generating an initial list of terms [28], and subsequently we searched the literature and consulted expert opinion to create a final list of 20 terms covering as much as possible the relevant behavioural expressions appropriate for the assessment of dogs in kennels. Eleven observers then used this list to score kennelled dogs in 13 video clips, and PCA of these scores generated four Principal Components (PCs) explaining 70.9% of the total variance. PC1 was characterised by the terms ‘curious/attention-seeking/playful/excited/sociable’, describing the animals’ interest in their environment and their engagement with people and pen-mates. PC2 ranged from ‘comfortable/relaxed’ to ‘anxious/nervous/stressed’, a demeanour apparently indicating the dogs’ ability to cope with and be comfortable in the kennel environment. PC3 describes fearful demeanour summarised by the terms ‘fearful/hesitant/wary’, whilst PC4 is characterised by the terms ‘depressed/bored’, which indicate a negative yet more low energy mood than fear, anxious and stress. Reference to emotional states with positive and negative valence has previously been made in assessments of shelter dog welfare: Titulaer et al. (2014) for example used cognitive measures to assess the effect of long versus short term permanence in kennel, whereas Mendl et al (2010) demonstrated how shelter dogs suffering from separation anxiety were more likely to be experiencing a negative affective state. As discussed in the introduction, Walker et al (2016) used comparable QBA dimensions to describe the emotional state of dogs in kennel versus home environment and compare short versus long term housing. Nevertheless, most of the scientific literature on shelter dogs’ welfare has relied, so far, mainly on behavioural and physiological indicators [38–40].

These four dimensions of emotional expressivity address dynamic aspects of welfare including important subtle differentiations, such as that between relaxation and depression, or between emotionally positive and negative excitement (excited vs nervous). From a ‘whole-animal’ perspective, the aim of QBA is not to identify a minimal set of core descriptive terms, but to capture wider patterns of expression and their context through a larger range of terms. Three terms (i.e. reactive, alert and aggressive) did not load highly on any of the four extracted PCs. Looking at Figures 1 and 2 it appears these terms co-load with stress and anxiety on PC2, and group together on PC4 opposite to ‘bored/depressed’, thus appearing to mostly reflect a tense reactivity associated with a negative emotional state in dogs struggling to cope with the environment. However, terms such as ‘alert’ and ‘reactive’ do not in themselves have a strong negative connotation – animals in positive playful mood can also be alert and reactive, or even mildly aggressive, and this likely explains why these terms often do not necessarily load highly on either positive or negative ends of emotional dimensions. This, however, does not make them superfluous, they can still serve to support and specify patterns of positive and negative mood in dogs assessed in different situations. Overall, the four PCs identified by this study cover the four quadrants of emotional expression defined by valence and arousal axes which tend to be typical of dimensional models of affect [41], and as such should be expected to offer a comprehensive assessment tool of dog emotional expression.

An important element of any measurement tool is its reliability, i.e. that measures used provide consistent results when applied by different assessors [42]. Inter-observer reliability with regard to QBA has two aspects: agreement in qualitative ranking of animals on expressive dimensions, and agreement in mean value of animal scores on expressive dimensions [43]. Agreement on qualitative ranking in the present study was good for all four PCs, (W=0.60-0.80), which aligns with previous studies testing the reliability of QBA fixed lists in a range of species [25,44,45], though some recent studies indicate that in field conditions as opposed to video-based conditions, achieving good agreement on qualitative ranking of animals on QBA dimensions can be more difficult [45,46]. There was however a significant observer effect on the mean value of animal scores for all four PCs suggesting differences between observers in how they used VAS scales for scoring across most terms. Post-hoc analysis, however, revealed that most observers (60-70%) did not differ significantly in their mean score for the 4 PCs, with fewer than 4 observers diverging significantly from this mean. Some of these outliers were often the same, suggesting that these observers had overall different scoring habits across all terms. For PC2 particularly mean scores were quite homogeneous, in fact, there was only one outlier. When running the analysis without that observer, no significant difference between the remaining observers was found. The higher homogeneity between scores associated to PC2 could be due to PC2 being the only dimension for which high loading terms described a clear contrast between positive and negative expressions. For the other three dimensions the emphasis was on either a positive or a negative expression, providing less conceptual anchoring for observers to place their scores on the visual analogue scale, making variation between observer scores more likely. Other studies have reported variation in ‘scoring styles’ between observers (e.g. [43]), creating observer effects on mean animal scores on some PCs.

Such effects are to an important extent due to differences in how observers score the various individual descriptors that are part of a QBA fixed-term list. Most QBA studies report differences in the level of agreement with which various individual terms were applied by observers (e.g. [25]) In the present study 12 out of 20 individual descriptors were applied at a level of agreement lower than 0.60, while 3 terms (depressed, explorative, aggressive) fell below 0.50. This lack of agreement may to some extent be due to the relatively low prevalence of these expressions in the video clips, however depressed demeanour may generally be difficult to distinguish reliably from other low energy expressions such as relaxed and/or bored [29,47]. Research on QBA assessment in donkeys and goats addressing such scoring issues under field conditions, has shown that training observers, i.e. taking time to discuss the meaning of individual terms and to compare how these terms are used for scoring, significantly improves the reliability of such terms [25,43]. Thus, any future practical application of the QBA term list for shelter dog welfare proposed in this study should provide ample training (including field assessments), until consistently high agreement between observers can be reached.

In recent efforts to promote the expression of positive emotions in captive/domestic animals (e.g. through appropriate environment enrichment), welfare scientists have directed a lot of attention toward the validation of outcome (i.e. animal-based) measures that could be integrated into welfare assessment tools to ensure that standards of positive welfare are maximised [48]. QBA has been successfully applied to farm animal welfare assessment protocols [22] and it has the potential to provide a valuable measure of dogs’ behavioural expression in confinement. Effective interventions to minimise signs of poor welfare in shelter dogs should be based on the evaluation of dogs’ individual ability to cope and adapt to confinement in kennels. QBA focuses on the dynamic expressivity of behavioural demeanour, describing and quantifying the emotional connotation of this expressivity that we naturally perceive. The comprehensive range of qualitative descriptors developed in this study may function as an integrative screening tool of kennelled dogs’ welfare, with special attention to positive welfare. A low expression of certain descriptors might prompt the implementation of certain types of enrichment that may have been neglected. For example, it has been demonstrated that stress upon entering a rehoming shelter can be mitigated by short sessions of positive human interaction [5]. Hence, low overall scores on the descriptors characterising PC1 may raise awareness for the need of targeted socialisation programs on animals struggling to adapt to the novel kennel environment. The QBA pre-fixed list developed in this study could be used as a monitoring system to detect early warning signs of reduced welfare state in kennelled dogs as well as detect areas of improvement to ensure good quality of life.

## 5. CONCLUSIONS

The present study aligns with previous QBA studies on other animal species in finding mainly good inter-observer reliability for a fixed-list of QBA descriptors applied to video-based assessments of dogs living in a shelter environment. Agreement in ranking dogs on the four expressive dimensions was good, but a few observers produced significantly different mean scores for observed dogs on the four main PCs, indicating a need for training and alignment of observer ‘scoring styles’ [43]. The scientific community recognises that welfare is a complex multidimensional concept, and that no single indicator can be considered exhaustive to evaluate the welfare of animals. It will be preferable therefore to apply QBA in conjunction with other physiological, behavioural or health indicators for animal welfare [49]. The QBA scoring tool developed and tested in the present study can be integrated into existing welfare assessment protocols for shelter dogs, and strengthen the power of those protocols to assess and evaluate the animals’ experience [7].

## Acknowledgments

The authors wish to thank the group of experts that anonymously participated in this study as well as all the observers for their contribution.

